# Clathrin light chain diversity regulates membrane deformation in vitro and synaptic vesicle formation in vivo

**DOI:** 10.1101/815183

**Authors:** Lisa Redlingshöfer, Faye McLeod, Yu Chen, Marine D. Camus, Jemima J. Burden, Ernest Palomer, Kit Briant, Philip N. Dannhauser, Patricia C. Salinas, Frances M. Brodsky

## Abstract

Clathrin light chain (CLC) subunits in vertebrates are encoded by paralogous genes *CLTA* and *CLTB* and both gene products are alternatively spliced in neurons. To understand how this CLC diversity influences neuronal clathrin function, we characterised the biophysical properties of clathrin comprising individual CLC variants for correlation with neuronal phenotypes of mice lacking either CLC-encoding gene. CLC splice variants differentially influenced clathrin knee conformation within assemblies, and clathrin with neuronal CLC mixtures was more effective in membrane deformation than clathrin with single neuronal isoforms nCLCa or nCLCb. Correspondingly, electrophysiological recordings revealed that neurons from mice lacking nCLCa or nCLCb were both defective in synaptic vesicle replenishment. Mice with only nCLCb had a reduced synaptic vesicle pool and impaired neurotransmission compared to wild-type mice, while nCLCa-only mice had increased synaptic vesicle numbers, restoring normal neurotransmission. These findings highlight differences between the CLC isoforms and show that isoform mixing influences tissue-specific clathrin activity in neurons, which requires their functional balance.

**SIGNIFICANCE STATEMENT:** This study reveals that diversity of clathrin light chain (CLC) subunits alters clathrin properties and demonstrates that the two neuronal CLC subunits work together for optimal clathrin function in synaptic vesicle formation. Our findings establish a role for CLC diversity in synaptic transmission and illustrate how CLC variability expands the complexity of clathrin to serve tissue-specific functions.

## INTRODUCTION

Clathrin mediates vesicle formation from the plasma membrane and endosomal compartments (1). Recruited by cargo-recognising adaptor proteins, triskelion-shaped clathrin proteins assemble into polyhedral lattices to capture membrane-associated cargo and promote membrane bending into clathrin-coated vesicles (CCVs). Through sequestration of a variety of cargo, CCVs play fundamental roles in general cellular physiology including regulation of nutrient uptake and signalling, as well as in tissue-specific membrane traffic such as synaptic vesicle (SV) generation (2). This range of clathrin functions has been attributed to adaptor and accessory protein variation (3). However, functional diversity is also generated by variability of clathrin subunits. Vertebrates have two types of clathrin heavy chains with distinct functions (4). The major vertebrate clathrin is formed from clathrin heavy chain CHC17 (herein referred to as CHC) associated with clathrin light chains (CLCs), which do not bind the minor CHC22 clathrin isoform. Vertebrate CLCs are encoded by two different genes *CLTA* and *CLTB* (in humans), producing CLCa and CLCb isoforms of about 60% sequence identity (5, 6). Their sequence differences have been conserved during evolution, after the encoding genes arose through duplication, suggesting the two isoforms can mediate distinct functions (7, 8). Expression levels of the CLCa and CLCb isoforms are tissue-specific (9), and further variation is created by alternative gene splicing during development (10) and in brain (5, 6). Here we address how CLC diversity affects the biophysical properties of clathrin and how the resulting variation affects the specialised function of clathrin in synaptic vesicle (SV) replenishment.

Limited functional differences between CLCa and CLCb have been observed in cell culture with respect to clathrin dynamics (11), during focal adhesion formation (12), cancer cell migration (13) and in epithelial polarity (14), with mechanisms attributed to isoform-specific differences in CLC binding proteins and post-translational modification (15). Loss of CLCs in culture and *in vivo* affects CCV uptake of some, but not all cargo, possibly reflecting variability in mechanical demand for packaging different cargo into CCVs (9, 15, 16). The capacity for biophysical differences in clathrin comprising different CLC isoforms and their splicing variants to influence tissue-specific clathrin functions has not yet been considered. CLC variability has potential to affect clathrin function as CLCs interact with key domains of the clathrin triskelion that contribute to clathrin-mediated membrane bending. They bind the trimerisation domain (TxD) of the triskelion vertex where the three component CHCs interact (17, 18), and extend along the triskelion leg to the bend at the knee where CLCs regulate conformation (19). Like various other endocytic proteins (3), CLCs undergo neuron-specific splicing (20), which introduces one exon in CLCb (encoding 18 residues) and two exons (encoding 30 residues) in CLCa at equivalent positions near the CLC C-termini, resulting in higher molecular weight forms nCLCb and nCLCa (5, 6), suggesting neuron-specific functions for these variants. *In vitro*, CLCs are required for efficient clathrin-mediated membrane vesiculation at low temperature (21). Thus, CLCs could affect clathrin-mediated membrane deformation directly through their influence on clathrin lattice properties (21, 22), as well as indirectly through cellular interaction with proteins that influence the actin cytoskeleton (14, 23-25).

SV replenishment by membrane traffic following neuronal degranulation is critical for sustained neurotransmission. Endocytosis is required to regulate synaptic plasma membrane surface area and to retrieve SV proteins (26), and SVs are generated from the plasma membrane and endosome-like compartments (27). The essential sites of clathrin function in these pathways are debated (2, 28), but dysfunction of clathrin-associated endocytic proteins is associated with neurological defects observed in Parkinson’s disease (29), Alzheimer’s disease (3) and in numerous animal models in which such proteins have been genetically deleted (30-35).

Here, we assess how clathrin lattices formed with neuronal and non-neuronal CLC variants could differentially affect function by correlating their *in vitro* biophysical properties with *in vivo* neuronal phenotypes of mice lacking CLC-encoding genes. We found that CLC composition significantly influenced clathrin lattice properties and the ability to form vesicles. CLC splice variation impacted assembly curvature by regulating CHC knee conformation, resulting in decreased ability of individual neuronal splice variants to deform liposome membrane *in vitro*. This effect was ameliorated for lattices formed from a mixture of neuronal CLC isoforms. Mice lacking either CLCa or CLCb isoforms showed corresponding differences in SV replenishment and synaptic neurotransmission compared to their wild-type (WT) littermates expressing both isoforms, and to each other. Collectively, these functional differences between CLC isoforms observed *in vitro* and *in vivo* establish a role for CLC diversity in regulating clathrin function in neurons and suggest a critical role for isoform balance.

## RESULTS

### CLC splice variants differentially influence lattice curvature via the triskelion knee

Within clathrin lattices, the knee bending angle determines whether a hexagon (non-curvature inducing) or pentagon (curvature-inducing) is formed (36, 37). In addition, the characteristic pucker at the triskelion vertex and the crossing angle of interacting CHC legs can further modulate lattice curvature (38, 39) (Fig. 1a). This versatility of clathrin assemblies allows clathrin to sequester cargo of various sizes as well as to form stable, flat assemblies that serve as signalling hubs (3, 40). For each closed cage, there is a fixed number of 12 pentagons, but a varying number of hexagons (41) (Fig. 1a). Thus, larger cages (i.e. lattices of lower curvature) have a smaller pentagon to hexagon ratio. CLCs maintain the puckered conformation of triskelia in flat assemblies (21) and structural studies showed that nCLCb can influence conformation of the triskelion knee up to a degree that inhibits assembly (19). Considering these effects on CHCs, we hypothesized that sequence differences between the CLC isoforms and alterations due to splicing might modulate CLC influence on triskelion conformation and lattice curvature.

**Fig. 1:**
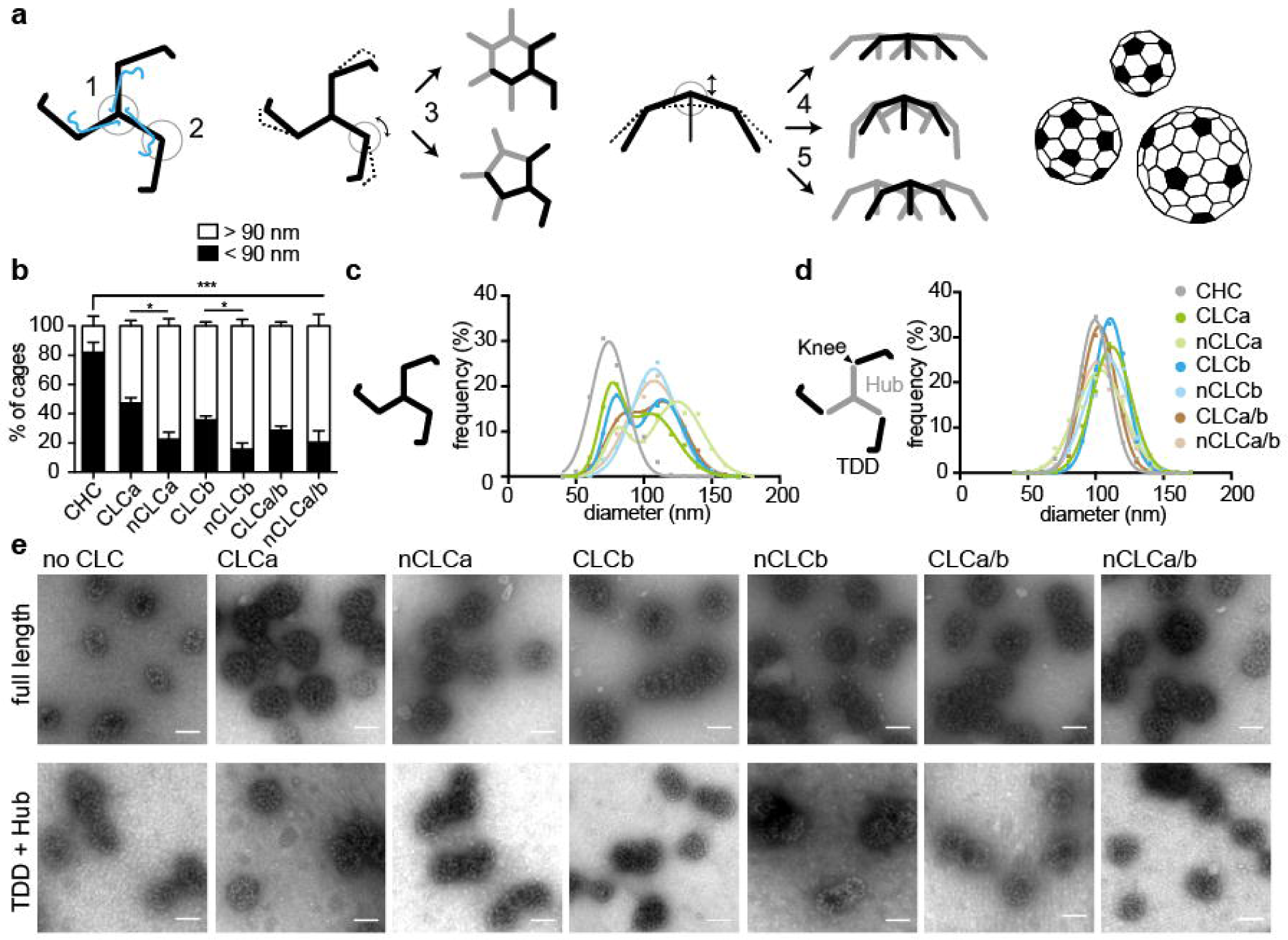
CLCs differentially affect lattice curvature via regulation at the CHC knee. **a** Pucker (1) and knee (2) angles of the clathrin triskelion (black: CHC, blue: CLC) dictate lattice architecture. Different knee angles (encircled, black straight and dashed lines) are adopted for hexagon (non-curvature inducing) or pentagon (curvature inducing) formation (3, whole triskelion in black and parts of others in grey). Lattice curvature is further amended through changes in pucker angle (4, dashed and black lines), or changes in proximal leg-crossing angle (5). CLC subunits are omitted for simplicity. Different sizes of closed cages are achieved by varying numbers of hexagons and a fixed number of 12 pentagons (filled black, based on (45)). **b** Quantification of the percentage of small cages (< 90 nm in diameter) in a population of more than 200 cages, generated by *in vitro* assembly of clathrin reconstituted with indicated CLCs, 1:1 mixtures of reconstituted clathrins (CLCa/b or nCLCa/b) or CHC only (CHC), determined from electron micrographs (mean ± SEM, \**P* < 0.05, \*\*\**P* < 0.001 one-way ANOVA followed by Holm-Sidak correction for multiple comparison, n = 3). **c** Cage size distributions from assemblies of triskelia formed from full length CHC as described in **b**, with CLC composition key in d. **d** Size distributions of cages formed from TDD (black) and Hub (grey) co-assemblies, with Hub fragments reconstituted or not with the indicated CLC isoforms or 1:1 mixtures thereof. **e** Representative EM images of cages formed from full-length CHC (top) or Hub/TDD fragments (bottom) without or following reconstitution with indicated CLC isoforms. Scale bar: 100 nm.

To address this, we produced clathrin triskelia of defined CLC composition by reconstituting tissue-derived CHC triskelia (42) with recombinantly expressed CLCa, CLCb, nCLCa or nCLCb (SI Appendix, Fig. S1a-d). We then induced assembly of these different clathrins into closed cages and analysed cage diameter as a measure of lattice curvature by electron microscopy. Previous studies showed that tissue-derived, CLC-bound clathrin (native) forms two major size classes, while CHC-only triskelia predominantly form small-sized cages (43-46). Our results confirmed that without CLCs, CHC formed predominantly cages of less than 90 nm in diameter with an average size of around 70 nm, representing cages of up to 60 triskelia (45). Clathrin with each of the CLC variants was further able to form larger cages with an average diameter of around 110 nm (140 triskelia) for those cages. The degree to which larger cages were formed varied significantly between splice variants (Fig. 1b, c), indicating that the neuronal splice inserts regulate the influence of CLCs on lattice curvature.

To establish whether differences in larger cage formation resulted from CLC influence on the triskelion vertex or on the CHC knee conformation or both, we dissected the effect of CLCs separately for each domain. To test whether CLC diversity exerts influence through varying CHC knee conformation, we produced clathrin cages from two CHC fragments that together constitute a full-length clathrin triskelion “cut” at the knee (Fig. 1d) (47). The Hub fragment (residues 1074-1675) was reconstituted with different CLCs (48) and combined with the terminal-distal leg segment (TDD, residues 1-1074, SI Appendix, Fig. S1e) under conditions promoting their co-assembly (47). Cages produced when TDD was combined with CLC-reconstituted Hub fragments were similar in size to fragment cages without CLCs, and the larger cages observed for CLC variants associated with intact CHC did not form (Fig. 1d, e). This result demonstrates that for CLC diversity to exert an effect on lattice curvature, the CHC knee must be intact, and shows that CLC splice variants differ in their influence on CHC knee conformation. Formation of larger cages by neuronal CLCs suggests that the neuronal splice inserts restrict the conformational flexibility of the CHC knee and could thereby reduce the likelihood of pentagon over hexagon formation. Non-neuronal CLCs appear to support a less biased variety of knee conformations, as cages of the two size classes are observed at almost equal frequency (Fig. 1b, c). To establish whether cage size differences also correlated with CLC effects at the TxD, the different CLCs were compared for their ability to stabilise the TxD (SI Appendix, Fig. S2) in an assay that measures triskelion dissociation. While dissociation differences were observed between triskelia reconstituted with the different CLC isoforms and their splice variants, these effects did not correlate with cage size differences, suggesting that CLC influence at the vertex is not a major factor in lattice curvature.

### CLC diversity modulates mechanical properties of the clathrin lattice

We next investigated whether the influence of CLCs on the CHC knee had mechanical consequences for their ability to support membrane deformation. Mechanical properties of clathrin lattices were previously shown to be CLC-dependent and reduced ability of CLC-free clathrin to bend membranes *in vitro* was correlated with reduced planar lattice quality when assembled on an electron microscopy (EM) grid (21). To establish whether CLC diversity influences lattice quality in the same assay, clathrins with different CLC composition were assembled on EM grids coated with a clathrin-binding fragment of epsin1 (H_6_-ΔENTH-epsin^144-575^, SI Appendix, Fig. S1f) (49). The flat lattices formed were visualized by EM and their periodicity (quality) assessed by Fourier transform analysis. Lattice quality was significantly reduced for clathrin with the neuronal splice variants of either CLCa or CLCb compared to clathrin reconstituted with their respective non-neuronal variants, and was significantly improved if formed from a 1:1 mix of nCLCa clathrin and nCLCb clathrin (Fig. 2). To visualise the nature of CLC distribution in these mixtures, lattices were generated using His-tagged nCLCa clathrin or nCLCb clathrin combined with the cognate untagged CLC and labelled with Ni-NTA-gold (SI Appendix, Fig. S3), confirming a uniform distribution of each neuronal CLC within the mixed lattice. In contrast to the behaviour of the neuronal CLCs, lattice quality was reduced in mixtures of CLCa clathrin and CLCb clathrin compared to lattices formed by clathrin with only CLCa or CLCb.

**Fig. 2:**
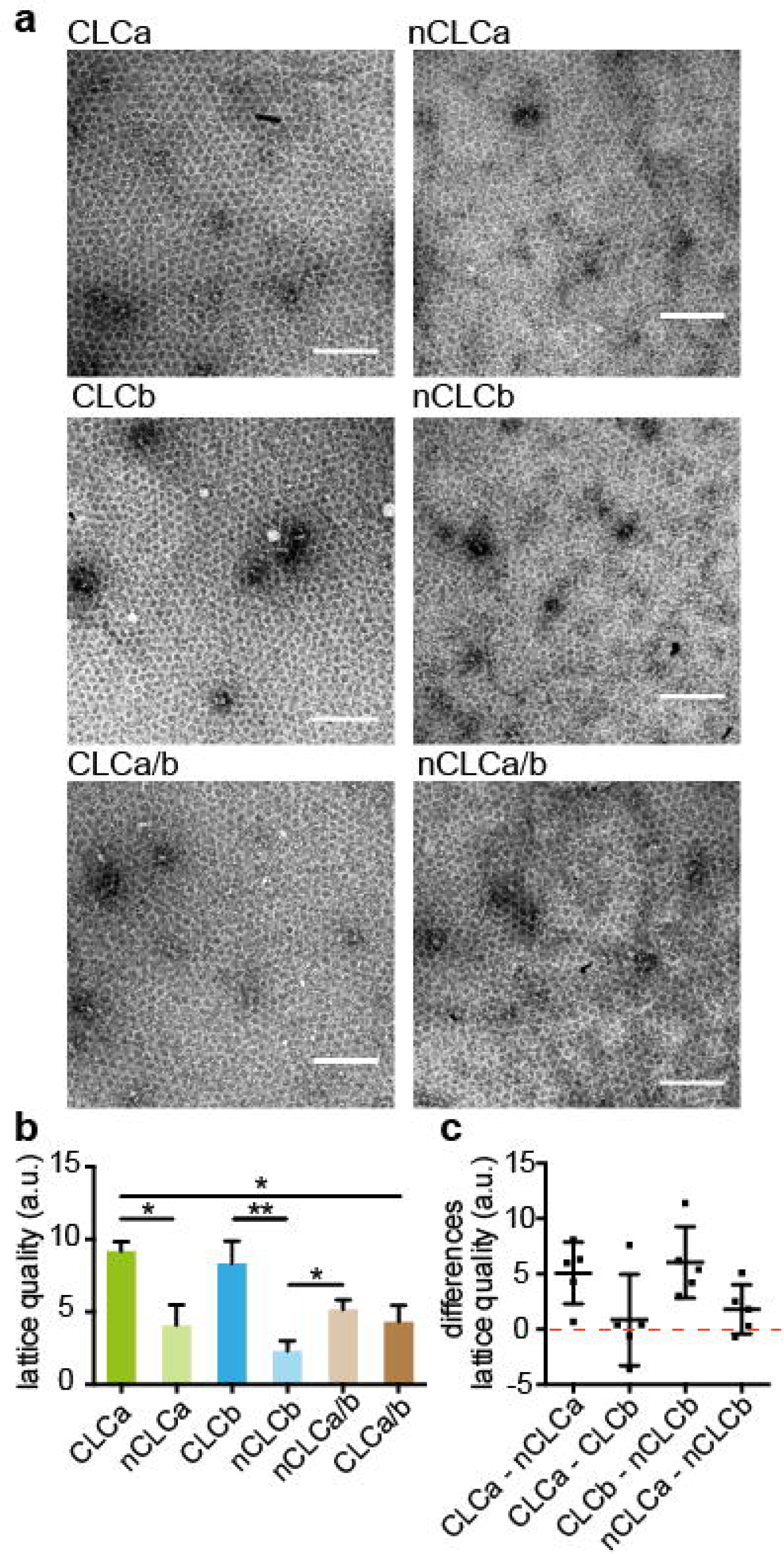
CLC splicing and mixing affects *in vitro* lattice quality. **a** Clathrin reconstituted with indicated CLC isoforms and mixtures thereof were assembled into flat lattices and visualised by negative stain for EM analysis. Scale bar: 200 nm. **b** Quality (regularity quantified by Fourier transform) of lattices generated as in **a**, (mean ± SEM, \**P* < 0.05, \*\**P* < 0.01, one-way ANOVA followed by Holm-Sidak correction for multiple comparison, n = 5). **c** Differences in lattice quality between indicated reconstituted clathrins in individual experiments as in **b** (data points and mean ± SEM, n = 5).

To determine whether CLC-related variation in lattice quality correlated with membrane bending efficacy, as seen for complete absence of CLCs, we tested the function of the CLC-reconstituted clathrins using a low temperature *in vitro* budding assay (21). In this assay, H_6_-ΔENTH-epsin^144-575^, coupled to liposomes via modified Ni-NTA lipids, captures clathrin at the liposome surface. Lattice assembly on these liposomes can generate mature buds, which remain attached to the liposome due to lack of dynamin, needed for scission (21, 49). The efficiency of clathrin with different CLC composition to form such coated buds was assessed at 15°C, a temperature permissive for budding by native clathrin but reduced for CLC-free clathrin (21). For each reconstituted clathrin, we measured the diameter of clathrin-coated membrane profiles in thin-section electron micrographs and assessed budding efficiency by the percentage of mature clathrin-coated buds (defined by a fitted diameter < 200 nm) relative to the total clathrin-coated membrane observed including shallow pits and flat clathrin assemblies (defined by a fitted diameter > 200 nm) (Fig. 3a). In line with previous findings, we found that at 15°C native clathrin was about twice as effective in membrane deformation as CLC-free clathrin (21), and that budding efficiency varied with the CLC composition of clathrin tested (Fig. 3b). CLCa or CLCb clathrin lattices generated mature membrane buds more efficiently than clathrin with either single neuronal clathrin variant (Fig. 3b, c), demonstrating that CLC splicing affects budding. Visualising the clathrin-coated liposomes absorbed to EM grids confirmed that all clathrins tested assembled into intact lattices or buds on the liposome surface (SI Appendix, Fig. S4), suggesting that the observed variation in budding was not due to impaired lattice formation, but CLC-dependent lattice properties (21). Notably, mixing reconstituted neuronal clathrins significantly improved budding efficiency compared to clathrins with only one type of neuronal CLC (Fig. 3d), overall reflecting the same pattern as observed for both lattice quality and curvature (Table 1), suggesting all three parameters result from differential CLC effects at the triskelion knee. Furthermore, all three assays demonstrated that neuronal clathrin benefits from cooperative co-assembly of clathrin with both isoforms.

**Table 1:** Regulation of clathrin lattice properties by CLC isoforms.

|  | high |  |  |  |  | low |
| --- | --- | --- | --- | --- | --- | --- |
| Lattice curvature | CLCb | CLCa | nCLCa/b | CLCa/b | nCLCa | nCLCb |
| Lattice quality | CLCb | CLCa | nCLCa/b | CLCa/b | nCLCa | nCLCb |
| Budding efficiency | CLCb | CLCa | nCLCa/b | CLCa/b | nCLCa | nCLCb |

**Fig. 3:**
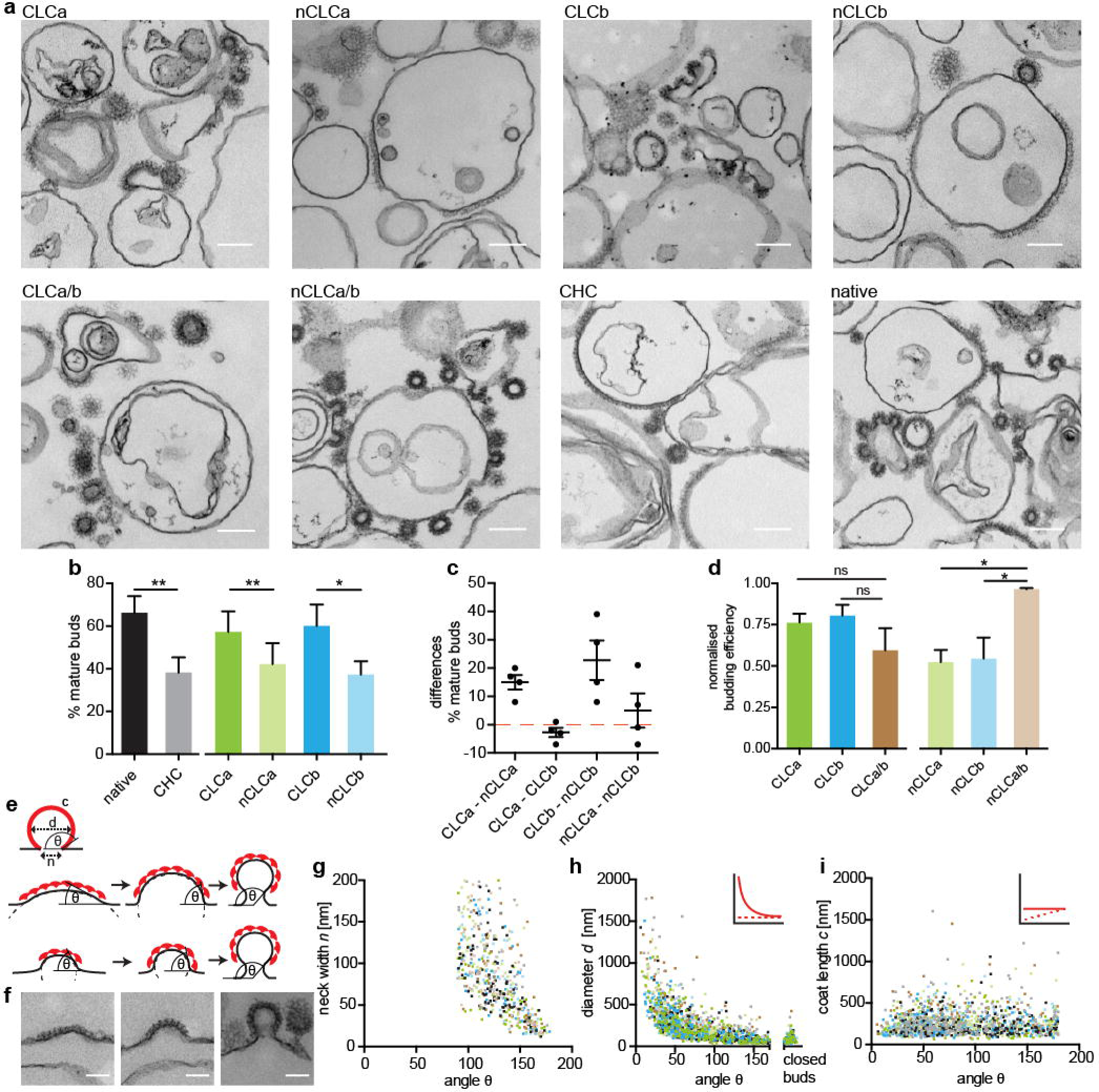
CLC splicing and mixing affects *in vitro* lattice budding efficiency. **a** Representative EM images of clathrin reconstituted with indicated CLC isoforms, 1:1 mixtures thereof, CHC only (CHC) and tissue-derived clathrin (native) assemblies on H_6_-ΔENTH-epsin^144-575^-coated liposomes. Scale bar: 200 nm. **b** Quantification of the percentage of mature buds (defined as clathrin-coated membrane profiles with < 200 nm fitted diameter) of more than 60,000 nm total clathrin-coated membrane profiles including flat, shallow and mature structures generated as in **a** (mean ± SEM, \**P* < 0.05, \*\**P* < 0.01, paired Student’s t-test for native and CHC, n = 3, one-way ANOVA with repeated measures for CLC-reconstituted clathrin, n = 4). **c** Differences in percentage of mature buds between indicated reconstituted clathrins in individual experiments in **b** (data points and mean ± SEM, n = 4). **d** Quantification of budding efficiency (% mature buds determined as in **b**, normalised to native) of reconstituted clathrins and mixtures thereof (\**P* < 0.05, one-way ANOVA for sets of neuronal or non-neuronal samples, followed by Holm-Sidak correction for multiple comparison, n = 3, p = 0.052, Student’s t-test for neuronal and non-neuronal mixtures, n = 3). **e** Parameters characterising coated buds (top); coat length *c*, membrane bud diameter *d*, neck width *n*, and budding angle *θ*. Models of clathrin-mediated membrane deformation by transitional curvature generation and lattice rearrangement (middle) or lattice growth under constant lattice curvature (bottom). **f** Representative coat profiles for flat (left), shallow (middle), and mature budded (< 200 nm fitted diameter, right) structures. **g** to **i** Analysis of the dataset generated as shown in **a** for coated membrane parameters *n* (**g**), *d* (**h**) and *c* (**i**) as shown in **e** in relation to the *θ* of each structure measured for all reconstituted clathrins within the same experiment. Inserts in **h** and **i** show the different correlations of these parameters as predicted according to the curvature transition (red, straight line) and constant curvature models (red, dashed line) shown in **e**.

Membrane deformation could result either from lattice formation at constant curvature or through transitioning from flat to curved lattices (Fig. 3e). Morphological analyses of clathrin-coated pits in human cell lines suggest that clathrin initially assembles into flat lattices at the plasma membrane and then gradually deforms the underlying membrane into coated buds (50, 51). To determine whether clathrin displayed these budding characteristics in our *in vitro* system and whether they are influenced by CLCs, we adopted the same EM-based approach. In micrographs of clathrin buds formed on liposomes, we measured the angle between the convex side of the coat and the coat-free membrane (Fig. 3e), coat curvature (i.e. diameter of the clathrin-coated structure), coat length and neck width for all coated membrane profiles (50) and then correlated these measurements. We observed a variety of coat curvatures (Fig. 3f) and found that neck width decreased with bud angle (Fig. 3g), as would be expected for both modes of deformation. Additionally, coat diameter correlated with the bud angle, characteristic of curvature transition (Fig. 3h), while we found no correlation between coat length and budding angle (Fig. 3i). Although this analysis is based on fixed structures at equilibrium, results were consistent with a flat to curved lattice transition, rather than assembly with constant curvature (Fig. 3h, i). Thus, we conclude that budding in our *in vitro* system displays the same characteristics as observed in cells (50, 51), where curvature is most likely generated gradually through lattice rearrangement. Collectively, our results suggest that CLC splice variants and mixtures thereof differ in their ability to promote this transition and that this is due to variable restrictive influence on the flexibility of the triskelion knee within the lattice, which in turn affects clathrin’s mechanical ability to deform membrane into coated buds (Table 1).

### CLC composition affects SV pool size and replenishment

Our *in vitro* experiments suggest that the CLCs differentially affect clathrin’s ability to deform membrane, with the neuronal isoforms working in conjunction for efficient membrane budding. Therefore, we predicted that neurons of animals with only one CLC isoform might be defective in clathrin-mediated pathways that rely on SV formation. This hypothesis was investigated in knock-out (KO) mice lacking the *Clta* or *Cltb* genes in all tissues (CLCa KO and CLCb KO mice). We previously generated CLCa KO mice (9) and have now produced CLCb KO mice. Loss of *Cltb* in CLCb KO mice was confirmed by PCR, and no CLCb or nCLCb protein was detected in all tissues analysed (SI Appendix, Fig. S5). Whereas wild-type (WT) mice express a mixture of CLCs in most tissues (9), CLCa KO mice express only CLCb or nCLCb, and CLCb KO mice express only CLCa or nCLCa, enabling functional analysis of clathrin with only one type of CLC in neurons (Fig. 4a).

**Fig. 4:**
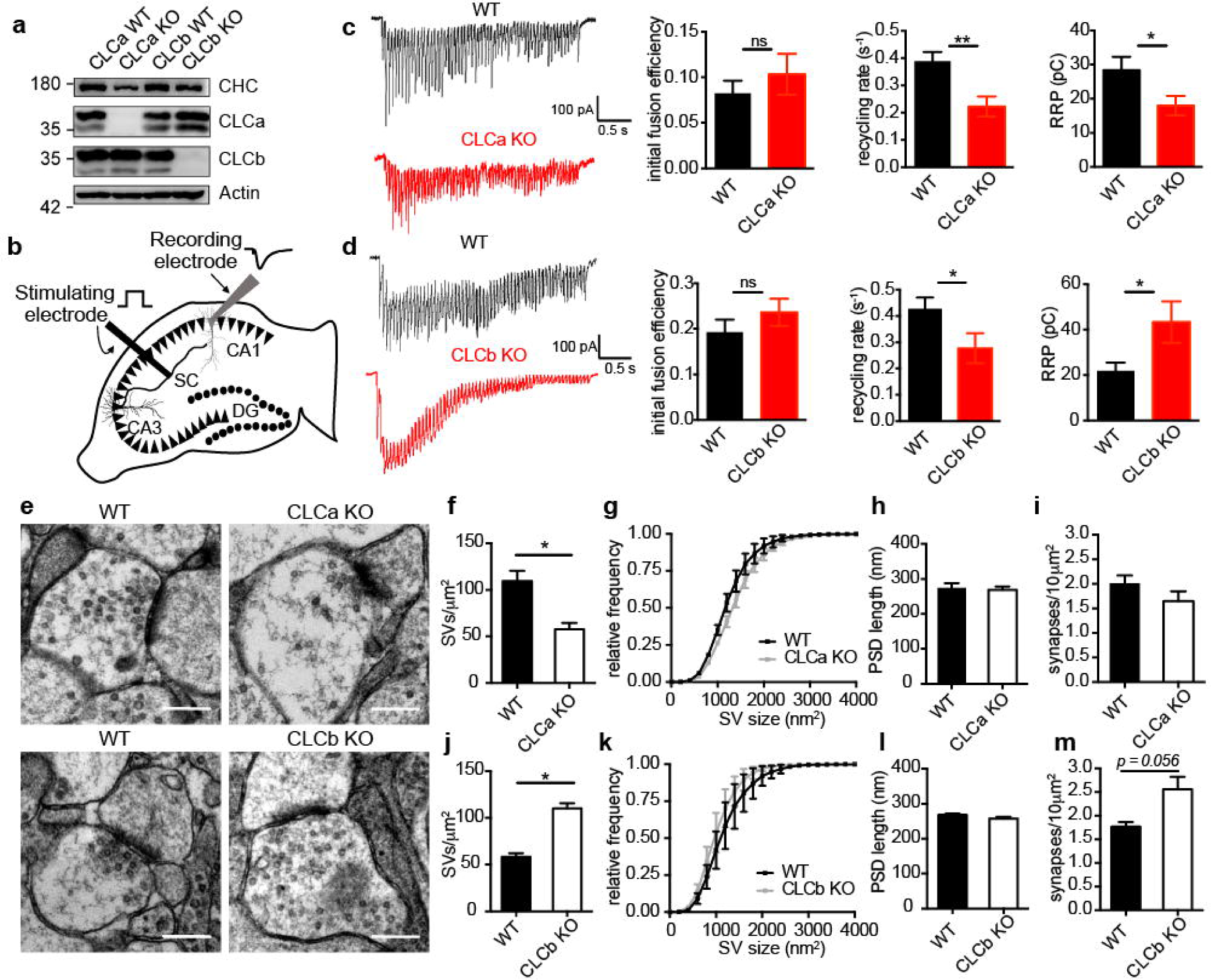
CLC composition regulates SV pool replenishment in hippocampal neurons. **a** Immunoblot of CLC KO and WT brain lysate for CHC (TD.1 antibody), CLCa (X16 antibody), CLCb (CLTB antibody) and actin (anti-beta-actin antibody). **b** Schematic illustration of stimulating and recording electrode setup in acute hippocampal slices. The synapses investigated are formed by a Schaffer-collateral (SC) axon (black) from a pyramidal neuron (black triangles, cell bodies) in the CA3 region synapsing with a receiving pyramidal neuron (grey triangle, cell body) in the CA1 region from which responses are recorded. Black circles denote dentate gyrus (DG) granule cell bodies. **c** Representative traces of evoked excitatory postsynaptic currents (EPSCs) following a 20 Hz electrical stimulation for 3 s in CLCa KO and WT hippocampal slices (left). Graphs (right) show the mean (± SEM) initial fusion efficiency, SV recycling rate and readily releasable pool (RRP) size calculated for all cells (n = 9-11 cells from 3 animals/genotype; \**P* < 0.05 and \*\**P* < 0.01, unpaired Student’s t-test). **d** Representative traces of EPSCs following a 20 Hz electrical stimulation for 3 s in CLCb KO and WT hippocampus slices (left). Graphs (right) show the mean (± SEM) initial fusion efficiency, SV recycling rate and RRP size calculated for all cells (n = 11-12 cells from 4 animals/genotype; \**P* < 0.05, unpaired Student’s t-test)**. e** Representative EM images of excitatory synapses in the CA1 region of the hippocampus of CLCa WT, CLCa KO, CLCb WT and CLCb KO. Scale bars; 300 nm. **f** to **m** Quantification of data extracted from EM images as in **e**. Graphs show SV density within 300 nm of the PSD (**e** and **i**), cumulative frequency distribution of SV size (**g** and **k**), postsynaptic density (PSD) length (**h** and **l**) and synapse density (**i** and **m**) expressed as mean ± SEM (\**P* < 0.05, unpaired Student’s t-test, n = 3).

Hippocampal neurons have frequently been used to study the function of endocytic proteins in synaptic transmission (27, 30, 31, 33, 34, 52). H&E staining of the hippocampus showed no morphological abnormalities in either KO strain, indicating that absence of either CLC isoform does not affect gross hippocampus development in the CLC KO mice (SI Appendix, Fig. S6). We therefore analysed the adult hippocampus in the CLC KO and WT mice to evaluate the function of mature excitatory synapses in the CA1 region, a region of well-defined neuronal architecture and neurophysiological circuitry. For electrophysiological recordings at Schaffer collateral (SC)-CA1 synapses in acute hippocampal slices, a stimulating electrode was placed in the SC fibres of the CA3 region and responses were then recorded in the CA1 pyramidal cell layer (Fig. 4b).

To determine the role of CLC diversity in SV pool replenishment *in vivo*, we recorded the responses to a prolonged high-frequency stimulus, which maximally depletes presynaptic terminals of the readily releasable pool (RRP) of SVs (53-55). Under these conditions, initial responses would draw from the pre-existing SV pool, while sustained neurotransmission would rely on the efficiency of SV pool replenishment (54). Using this approach, we were able to assess CLC-dependency for different stages of SV exo- and endocytosis from these recordings. We found that the initial fusion efficiency of SVs in either KO strain was similar to that of their WT littermates, indicating that packaging of fusion-mediating cargo was not affected by changes in CLC composition (Fig. 4c, d and SI Appendix, Fig. S7). Instead, both KO models had a decreased SV recycling rate characteristic of a defect in acute SV pool replenishment (Fig. 4c, d and SI Appendix, Fig. S7). This finding correlated with our expectation from the *in vitro* studies that showed clathrin lattices with only one type of neuronal CLC were different from mixed lattices in their assembly properties (Table 1).

Further calculations based on data obtained from sustained trains of action potentials indicated that the RRP was larger in the CLCb KO mice but reduced in the CLCa KO mice when compared to WT littermates (Fig. 4c, d and SI Appendix, Fig. S7), revealing a phenotype that we did not expect from our *in vitro* data. To establish whether the SV formation was altered in the CLC KO mice rather than the ability to mobilise SVs from other SV pools, we assessed the ultrastructure of excitatory synapses in the CLC KO mice and their WT littermates (Fig. 4e). Indeed, SV density was significantly reduced in hippocampal neurons in the CLCa KO animals. In contrast, SV density in equivalent neurons of the CLCb KO mice was significantly increased compared to their WT littermates (Fig. 4f, j), in line with our RRP estimates from electrophysiological recordings (Fig. 4c, d). Other parameters such as individual SV size (Fig. 4g, k), postsynaptic density (PSD) length (Fig. 4h, l) and excitatory synapse density (Fig. 4i, m) were similar between both KO strains and respective WT littermates. Together, these data show that CLCs are required for efficient and sustained synaptic function and suggest that loss of either CLC decreases acute SV replenishment by affecting immediate formation, correlating with expectations from reduced in vitro budding efficiency, but that the KO strains differ in their ability to generally maintain SV pools. Thus, CLC composition influences SV density in excitatory neurons of the hippocampus, and nCLCa and nCLCb function differently in SV generation such that mice with only nCLCa can sustain a compensatory pathway for SV generation and mice with only nCLCb cannot.

### CLCa KO and CLCb KO mice have distinct neurological and behavioural defects

To further investigate the differential consequences of CLCa or CLCb KO for synapse function, we conducted additional electrophysiological experiments to characterise synaptic transmission in our KO animals. Analyses of the evoked excitatory postsynaptic current (EPSC) responses to stimuli of increasing magnitude revealed decreased EPSC response amplitude in the CLCa KO mice compared to WT littermates, suggesting impaired basal synaptic transmission in the CLCa KO mice (Fig. 5a). In contrast, the evoked EPSC response amplitude in hippocampal slices from the CLCb KO mice was of similar or higher magnitude than that of their WT littermates, suggesting that basal synaptic transmission was intact and even enhanced (Fig. 5b). These differences in synaptic connectivity between hippocampal function in the two KO strains could arise from defects in presynaptic neurotransmitter release correlating with differences in SV pool size in the KO animals relative to their WT littermates (Fig. 4).

**Fig. 5:**
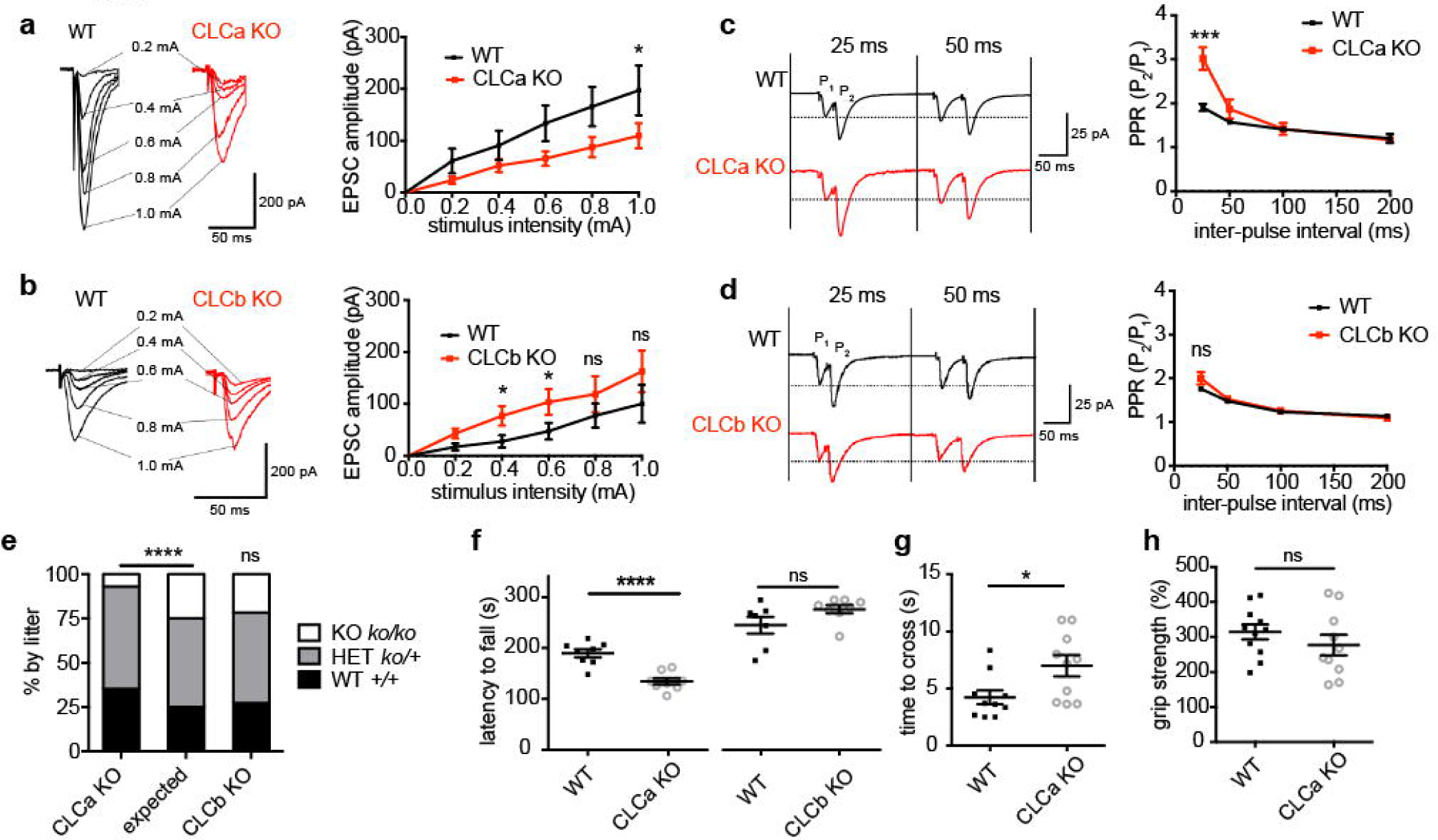
Neuronal defects are compensated for in CLCb KO but not CLCa KO mice. **a** Input-output relationship of evoked excitatory postsynaptic currents (EPSCs) at CA1 synapses of WT and CLCa KO hippocampus slices. Traces show responses of representative cells at increasing stimulus intensity with an average of 3 responses for each stimulus strength (mean ± SEM; n = 9-11 cells from 3 animals/genotype; \**P* < 0.05, two-way ANOVA with repeated measures). **b** Input-output relationship of evoked EPSCs at CA1 synapses of WT and CLCb KO hippocampus slices. Traces show response of representative cells at increasing stimulus intensity with an average of 3 responses for each stimulus strength (mean ± SEM; n = 12-15 cells from 4 animals/genotype; \**P* < 0.05, two-way ANOVA with repeated measures). **c** Paired-pulse ratio (PPR) of evoked EPSCs at CA1 synapses of WT and CLCa KO hippocampal slices. Traces show responses of representative cells. Graph displays the mean PPR (P_2_/P_1_ ± SEM) from all cells (n = 9-11 cells from 3 animals/genotype) at different time intervals between paired pulses (\*\*\**P* < 0.001, unpaired Student’s t-test). **d** PPR of EPSCs at CA1 synapses of WT and CLCb KO hippocampal slices. Traces show responses of representative cells. Graph displays the mean PPR (P_2_/P_1_ ± SEM) from all cells (n = 12-15 cells from 4 animals/genotype) at different time intervals between paired pulses. **e** Genotype distribution for CLCa KO (n = 590) and CLCb KO (n = 359) mice after weaning compared to expected distribution (\*\*\*\**P* < 0.0001, *Chi*-square test). **f** Performance in the accelerated rotarod test (latency to fall) by CLC KO (CLCa KO = 8, CLCb KO = 8) and control wild-type littermates (CLCa WT = 8, CLCb WT = 7) expressed as data points and mean ± SEM (\*\*\*\**P* < 0.0001, unpaired Student’s t-test). **g** Time to cross grid for CLCa KO (n = 10) and control mice (n = 10) expressed as data points and mean ± SEM (\**P* < 0.05, unpaired Student’s t-test). **h** Grip strength of CLCa KO (n = 10) and control mice (n = 11, gram-force relative grip strength over body weight) expressed as data points and mean ± SEM.

We then performed paired-pulse ratio (PPR) recordings experiments, an approach that allows assessment of defects in neurotransmitter release (56, 57). In excitatory neurons such as those analysed here, a second stimulus pulse (P_2_) fired in short succession after an initial pulse (P_1_) results in a larger response than the first stimulus. This facilitation is dependent on changes in the probability of neurotransmitter release. PPR is inversely correlated with release probability, as a lower initial release probability leaves more vesicles remaining at the terminal which are then capable of being released after the second stimulus (56, 57). We observed that, compared to WT littermates, the PPR was increased in CLCa KO mice, consistent with a reduction in release probability (Fig. 5c). In contrast, no differences in PPR were observed in CLCb KO mice compared to WT littermates (Fig. 5d). Thus, specific loss of nCLCa impairs presynaptic function compromising neurotransmitter release at the SC-CA1 synapse. Collectively, our electrophysiological data support the morphological data (Fig. 4e-m), indicating that neurons with only nCLCa clathrin can compensate for defective SV regeneration by expanding their SV pool from another pathway, while neurons with only nCLCb clathrin cannot compensate and show decimated SV pools from impaired replenishment.

Ongoing breeding of the CLCa KO colony confirmed a 50% survival rate compared to that expected for homozygous CLCa KO mice (9), whereas homozygous CLCb KO mice had no survival defects (Fig. 5e). Survival phenotypes similar to the CLCa KO animals have been reported for other genetic deletions in endocytic pathways, which also displayed neuronal phenotypes (30) in the hippocampus and other regions of the brain (32). Therefore, we assessed performance in the rotarod test for neuro-motor coordination for both CLC KO mice (58). Compared to their WT littermates, surviving CLCa KO mice displayed defects in rotarod balance, whereas CLCb KO mice did not (Fig. 5f). Further assessment of sensorimotor function revealed that CLCa KO mice exhibited defects in a grid-walking test (Fig. 5g), but not in grip strength (Fig. 5h), suggesting neurological rather than muscular dysfunction in the CLCa KO animals, supporting the notion that loss of nCLCa is less tolerable than loss of nCLCb. Together, the phenotypes of the KO animals indicate that neuronal CLC isoforms are differentially able to support compensatory mechanisms for adjusting SV pools hippocampal synapses with impaired acute SV replenishment, and suggest that clathrin with only nCLCa enables more functions than clathrin with only nCLCb, while a mix of both neuronal CLCs is necessary for optimal clathrin function in neurons, in keeping with their synergistic contribution to clathrin function *in vitro.*

## DISCUSSION

To understand the consequences of CLC diversity for clathrin function, we characterised the biophysical properties of *in vitro* assemblies formed from clathrin comprising single CLC isoforms and correlated these properties with neuronal phenotypes observed in KO animals expressing single CLC isoforms. We found that neuronal CLC splicing affects lattice properties (Fig. 1b and Fig. 2b) by diversifying the CLC influence on the CHC knee to regulate lattice curvature (Fig. 1c, d) and deform membrane (Fig. 3b). Our *in vitro* studies further suggest that lattices formed from mixtures of clathrin with nCLCa and nCLCb have different assembly properties and are more efficient in membrane deformation compared to clathrin with only one type of neuronal CLC (Fig. 2b, 3d). Consistent with this observation, neurons from both CLCa and CLCb KO mice showed defects in synaptic transmission that indicated acute clathrin-dependent SV replenishment was impaired (Fig. 4c, d). However, the CLCa KO mice had reduced numbers of SVs in their hippocampal synapses, while the CLCb KO mice had more SVs than their WT littermates (Fig. 4f, j). Thus, although acute SV replenishment was impaired in both KO strains, clathrin with only nCLCa (in the CLCb KO mice) was able to support a compensatory pathway of SV formation, while clathrin with only nCLCb (in the CLCa KO) was not (Fig. 5a-d). These findings establish functional differences between CLC isoforms *in vitro* and *in vivo* and demonstrate how resulting clathrin diversity, as well as CLC isoform balance, are important for clathrin function in neurons.

Neuronal splicing predominates for CLCb in neurons (59), while neuronal CLCs are not present in other brain cells such as glial or Schwann cells (20), indicating that the CLC splice variants segregate within brain cell types to fulfil distinct functions. Neuronal CLC variation due to splicing had a significant influence on lattice curvature and budding efficiency. Compared to clathrin with non-neuronal CLCs, clathrin with neuronal CLCs formed predominantly large cages in solution (with low a pentagon to hexagon ratio), suggesting that through their influence on the CHC knee (Fig 1d), the spliced inserts reduce triskelion flexibility to accommodate pentagonal faces. When constrained to a flat, solid surface (EM grid), lattice assembly requires significant deformation of triskelia (21). The quality of lattices formed by clathrin with neuronal CLCs under these conditions was reduced compared to clathrin with non-neuronal CLCs, further suggesting that neuronal CLCs restrict clathrin’s conformational flexibility. Notably, CLC-dependent lattice quality and cage size correlated with the ability of clathrin with different CLCs to produce mature buds at liposome membranes, which morphological analysis suggested are generated by flat lattice rearrangement, as observed at cell membranes (50, 51). Together, these results suggest that CLC splicing variation influences clathrin’s flexibility to form and alternate between a range of morphologies in order to deform lipid membrane into mature clathrin-coated buds, and that this is attributable to CLC influence on the CHC knee domain.

Given that the inserted sequences are located near the TxD where the C-termini of CLCs are bound (arrowheads, Fig. 6a), we propose that the splice inserts affect the conformation of the adjacent knee of a neighbouring triskelion, which is closer to the inserted sequences than the knee of the triskelion to which the CLC is bound (Fig. 6b). This intermolecular influence could involve splice inserts at the C-terminus (stars, Fig. 6b) interacting with the neighbouring CHC knee or with the N-terminal domain (N) of a CLC bound to the neighbouring knee (Fig. 6b). Considering the average 40% sequence differences between CLCa and CLCb isoforms and their even greater variation at the N-terminus, inter-CLC interactions could vary depending on which splice isoform interacts with which CLC isoform N-terminus, thereby influencing overall lattice properties. This imputed interaction may be lost from clathrin with non-neuronal CLCs, which would explain why mixing non-neuronal CLC isoforms does not affect *in vitro* membrane deformation to the same extent as mixing neuronal CLC isoforms (Fig. 3d).

**Fig. 6:**
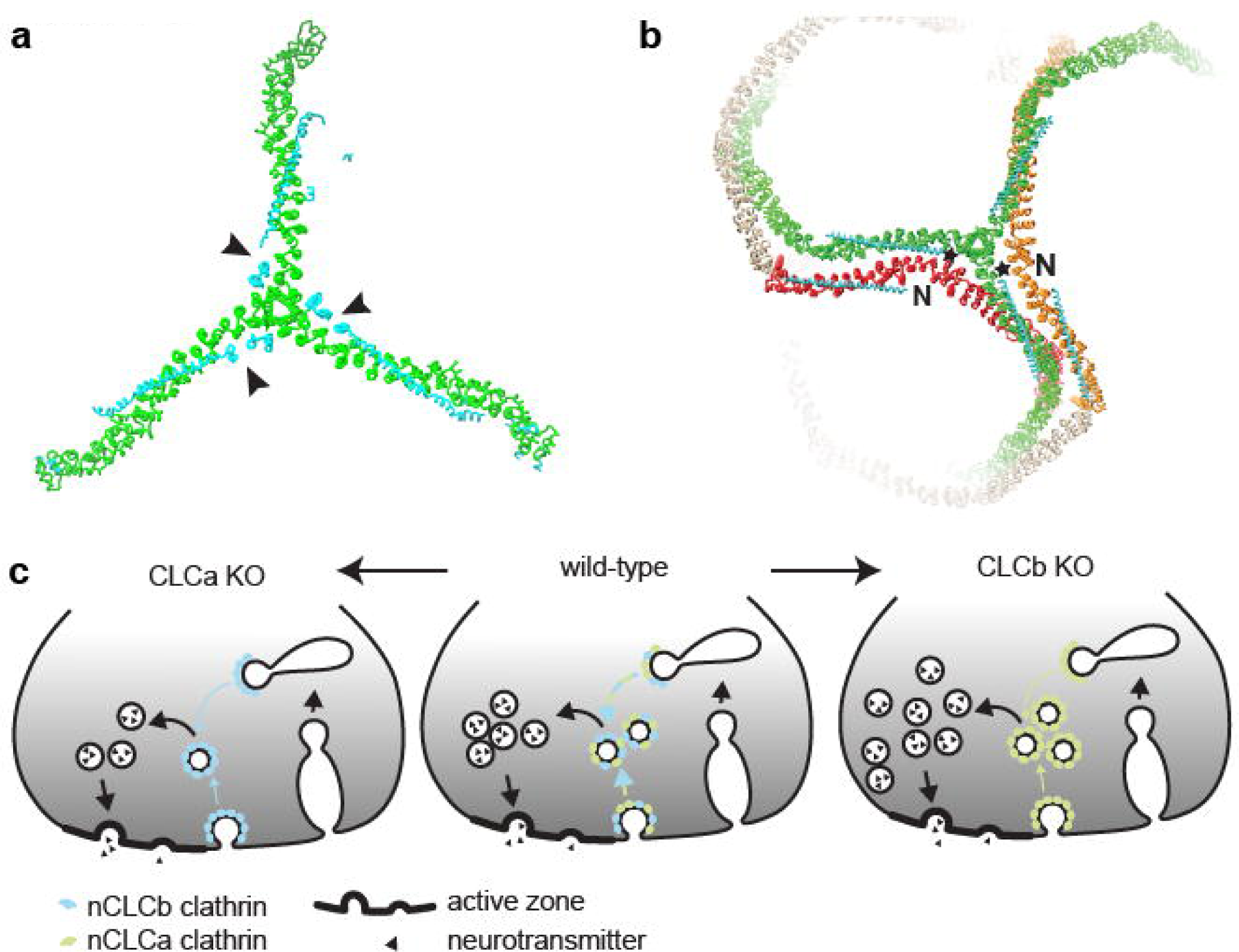
Model of how neuronal CLC diversity regulates synaptic vesicle recycling and lattice properties. **a** CLC (cyan) neuronal splice inserts (arrowheads) are located near the TxD of the bound CHC (green, left, PDB: 3LVH). **b** CLC (cyan) neuronal splice inserts (stars) are near the CHC knee of the neighbouring triskelion (red or orange) within lattices (right, PDB: 3IYV). Interaction between a neuronal splicing insert and the knee of an adjacent triskelion and/or adjacent CLC N-terminus (N) could promote conformational change in the CHC knee. As the formation of pentagons and hexagons requires different knee angles, this interaction could consequently influence lattice curvature (see Fig. 1a). **c** In WT mice (centre), a mix of nCLCa and nCLCb clathrin creates the appropriate biophysical properties to mediate SV generation from endosomal compartments (and possibly the plasma membrane, blue-green arrows). Loss of nCLCa (left) creates nCLCb clathrin lattices that are defective in efficient SV regeneration (blue arrows), resulting in decimated SV pools. Loss of nCLCb (right) creates nCLCa clathrin lattices, which are also less efficient in SV regeneration than WT, but able to maintain an overall increased SV pool by excess budding to compensate for reduced efficiency in acute SV pool replenishment (green arrows).

Here, we characterise the properties of homo- and hetero-assemblies of clathrin comprising single CLC isoforms. In brain, CLC isoforms are apparently randomly distributed on triskelia (60), in which case neuronal inserted sequences could still influence lattice curvature via interactions with an adjacent triskelion knee. The presence of nCLCb would serve as an attenuator of nCLCa-specific interactions and vice versa. Further, in cells where CLCb expression is transiently increased (61), newly synthesized triskelia would be generally occupied by a single CLC isoform as clathrin subunit turnover is slow (62). The resulting homotypic clathrins would then participate mixed in lattices, as we have studied here. Thus, neuronal CLC isoform expression ratios (and/or local abundance within cells) could tailor clathrin lattice properties specifically to particular needs.

There are several pathways involved in SV recycling in neurons and the role of clathrin has been widely debated, possibly because the contribution of each pathway to SV formation is variable between organisms, types of neurons and stimulus (52, 63, 64). SV recycling can either be achieved by direct regeneration of SVs through clathrin-mediated endocytosis from the plasma membrane at low-frequency stimulus, or through clathrin-independent mechanisms, which predominantly mediate retrieval of SV components from the plasma membrane under high stimulus (2, 27, 28, 33, 52). Rapid clathrin-dependent trafficking of SV proteins from the endosomal pathway then leads to re-sorting and formation of SVs at high-frequency stimulus (2), which can occur in a timeframe of 1-3 seconds after stimulation (27). In our analysis of neurotransmission, SV recycling defects were detected by electrophysiology within three seconds at high stimulus (20 Hz for 3s), suggesting the defects detected were due to impaired clathrin-mediated regeneration of SVs from the endosomal pathway. These findings are consistent with previously observed roles for CLCs in recycling from non-neuronal endosomal compartments (61, 65).

That neurons in both CLC KO strains shared an acute defect in SV pool replenishment fits with our *in vitro* biophysical data that demonstrate single neuronal CLCs are not as efficient in membrane deformation as a mixture of clathrin with both neuronal CLCs (Fig. 3d). In response to this defect, we found that clathrin with only nCLCa generated an increased steady-state SV pool in CLCb KO neurons, while clathrin with only nCLCb could not (Fig. 4). CLCa seems to have preferential connection with the actin cytoskeleton compared to CLCb (12, 14), which might account for the ability of nCLCa clathrin to function on its own without nCLCb and benefit from involvement of actin and its accessory molecules in this compensatory pathway. It is possible that in the (normal) presence of nCLCb, the actin interactions of nCLCa may be “diluted down”, so that nCLCb works as an attenuating balancing mechanism to control SV pools at steady state. The presence of nCLCb would simultaneously create lattice properties for efficient CCV budding during neurotransmission. In its absence, the acute replenishment pathway would decrease in efficiency but loss of the nCLCb attenuation effect enables a compensatory pathway with nCLCa only (Fig. 6c). Thus, the budding defect resulting from a change in biophysical properties of clathrin is detectable only under acute demand, and when demand is less acute, there are mechanisms by which nCLCa clathrin can generate SVs that are not supported by nCLCb clathrin.

Considering the whole animal phenotypes, including the increased mortality rate for CLCa but not CLCb KO animals, CLCa seems able to sustain house-keeping clathrin functions, while CLCb functions mainly in conjunction with CLCa, apparently in a regulatory capacity, such that a balance between the two is required for some specialised functions. This is in line with the observation that CLC expression ratios vary with tissue, and while lymphoid cells almost exclusively express CLCa, no tissue has been found to express exclusively CLCb (9, 62). CLC isoform splicing changes during cell differentiation, development (10, 59) and ratios of CLCb relative to CLCa are modulated in tumour progression (13) and cell migration (61). These observations, in combination with the *in vitro* and *in vivo* data reported here, support the concept that, in addition to influencing accessory protein interaction, the conserved CLC isoform and splicing differences characteristic of vertebrates facilitate tissue-specific clathrin functions by directly modulating clathrin lattice properties. In particular, CLC diversity regulates clathrin budding efficiency, a property that we show here affects SV formation under acute demand, and is likely to influence other pathways that rely on rapid clathrin-mediated membrane traffic.

## MATERIALS AND METHODS

### Protein expression and purification

Except for full-length CHC, all proteins were recombinantly expressed in bacteria and purified by standard affinity and size exclusion chromatography methods as specified in SI Appendix, Materials and Methods. Native clathrin and full length CHC were prepared from porcine brain as specified in SI Appendix, Materials and Methods.

### Characterisation of clathrin lattice properties

For lattice curvature determination, cage assembly was induced by dialysis in assembly-promoting buffer A (100 mM MES, 1 mM EGTA, 0.5 mM MgCl_2_, 2 mM CaCl_2_) and cage size distributions analysed form EM images as specified in SI Appendix, Materials and Methods. Flat lattices were produced and analysed as previously described (21) and as detailed in SI Appendix, Material and Methods. Lattice regularity was assessed from electron micrographs of similar quality and various focuses for each specimen as a measure of lattice quality. 5-12 images were analysed for each reconstituted clathrin isoform within each of three independent sets of experiments. *In vitro* budding was performed as previously described (49). In brief, H_6_-ΔENTH-epsin^144-575^ was bound to liposomes made from brain polar lipid extracts containing 5% DGS-Ni-NTA lipids (Avanti). H_6_-ΔENTH-epsin^144-575^-coated liposomes were then chilled to 15°C and mixed with clathrin triskelia at 15°C, incubated for 30 min at 15°C and then transferred to ice. Samples were then fixed at 4°C overnight and processed for EM analysis. Diameter, coat length, bud angle and neck width of clathrin-coated membrane profiles were measured from electron micrographs using ImageJ (NIH) and data processed using Prism (GraphPad). Between 150 and 200 coat profiles, randomly sampled across four thin sections, were analysed for each sample (> 60,000 nm total coat length per experiment per sample). The proportion of mature buds, defined by a diameter of less than 200 nm, of all clathrin-coated membrane profiles examined, was used as a measure of budding efficiency. Results from 3-5 independent sets of experiments were tested for statistical significance.

### Generation of CLC KO mice

The *Cltb*^*ko/ko*^ mouse strain was created from ES cell clone 19159A-F4, generated by Regeneron Pharmaceuticals, Inc., and obtained from the KOMP Repository (www.komp.org). Methods used to create the CLCb-null ES cell clone have previously been published (66). In brief, the complete coding region of the *Cltb* gene was fully deleted by homologous recombination using a large BAC-based targeting vector. Targeted ES cells were then injected into albino C57BL/6J-N blastocytes and transferred into foster mothers. Chimeric offspring were mated with C57BL/6J females (Charles River), and germ-line transmission of the *Cltb*-null allele (*Cltb*^*ko*^) was established. Heterozygote *Cltb*^ko/+^ mice were backcrossed on the C57BL/6 background and bred to produce *Cltb*^ko/ko^ homozygous mice. CLCa KO mice were derived from C57BL/6 WT mice (9) and produced by breeding *Clta*^ko/+^ heterozygotes. WT littermates from the same breedings were used as controls for homozyogous KO animals. All procedures involving animals were conducted according to the Animals Scientific Procedures Act UK (1986) and in compliance with the ethical standards at University College London (UCL).

### Analysis of protein expression in brain

Mouse brains were harvested from six 9-12-month old C57BL/6 WT and homozygous CLCa and CLCb KO mice and snap-frozen in liquid nitrogen and stored at −80°C until further use. Tissue were quickly thawed and homogenised in lysis buffer (50 mM HEPES pH 8.0, 50 mM NaCl, 1% Triton-X 100, 5 mM EDTA, 2 mM CaCl_2_, 1 mM PMSF, cOmplete™ Protease Inhibitor Cocktail mix (Roche)). The homogenate was further incubated on ice for 45 min before centrifugation at 21,000 g and 4°C for 2 x 10 min in an Eppendorf 5424 R benchtop centrifuge (Eppendorf). Protein content of the lysate was determined by BCA assay (Thermo Fisher). For analysis, 25 μg of sample was loaded on 4-15% acrylamide gels (Bio-Rad) and subjected to SDS-PAGE and immunoblotting. Primary antibodies used for immunoblotting were TD.1 (anti-CHC, made in house) (67), X16 (anti-CLCa, made in house) (68), CLTB (anti-CLCb, Proteintech) and anti-beta-actin (Sigma-Aldrich).

### Electrophysiology

Experiments were performed in 8-10-month-old mice. Both male and female mice were used for electrophysiological experiments. Acute transverse hippocampal slices (300 µm) of homozygous CLCa or CLCb KO and control mice were cut on a Leica VT-1000 vibratome in ice-cold artificial cerebrospinal fluid (ACSF) bubbled with 95% O_2_/5% CO_2_ containing (in mM): NaCl (125), KCl (2.4), NaHCO_3_ (26), NaH_2_PO_4_ (1.4), D-(+)-Glucose (20), CaCl_2_ (0.5) and MgCl_2_ (3). At 5-minute intervals, slices were then transferred into a series of 3 different chambers oxygenated (95% O_2_/5% CO_2_) in the same base ACSF but with the following temperature and component (in mM) variations: **1.** 21°C initially with MgCl_2_ (1) and CaCl_2_ (0.5) then allowed to heat gradually to 36°C; **2.** 36°C with MgCl_2_ (1) and CaCl_2_ (1); and **3**. 36°C initially with MgCl_2_ (1) and CaCl_2_ (2) before cooling to 21°C. Slices were then left for at least 1 hour before recordings commenced.

Evoked recordings were performed on an upright microscope continually perfused with oxygenated recording solution at room temperature containing the same ACSF composition as the third chamber and supplemented with 10 μM bicuculline. Pyramidal cells in the CA1 region were held at −60 mV in whole-cell voltage-clamp configuration using glass microelectrodes (resistance 3-8 MΩ) filled with caesium gluconate intracellular solution containing (in mM): D-gluconic acid lactone (130), HEPES (10), EGTA (10), NaCl (10), CaCl_2_ (0.5), MgCl_2_(1), ATP (1) and GTP (0.5), QX314 (5), pH to 7.2 with CsOH. To evoke postsynaptic EPSCs, a bipolar concentric stimulation electrode (FHC) was placed in the SC fibres of the CA3 region. For input-output recordings, the stimulus pulse was varied between 0.2 and 1 mA with a pulse width of 0.1 millisecond (ms) and stimuli were delivered at a rate of 0.1 Hz. Paired pulse stimuli were given at rate of 0.2 Hz with different inter-stimulus intervals, ranging from 25 ms to 200 ms and a stimulation strength set to approximately 50% of the maximal response for each cell. PPR was calculated as the ratio of the peak amplitude of the second response over the first response. Calculation of RRP size, initial fusion efficiency and SV recycling rate was done on ESPC recordings that underwent 3 s duration trains of stimulation at 20Hz and estimated as previously described (54, 69). Briefly, RRP size, fusion efficiency (*fe*) and vesicle recycling rate (α) were evaluated from the cumulative charge during the stimulation train using the following two equations:

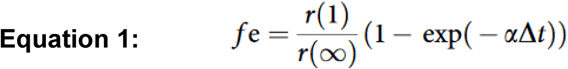

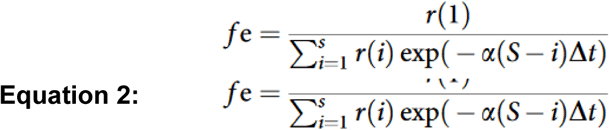

*r*(1) is the charge of the first EPSC in the train

*r*(i) the charge passed by the *i*th EPSC

*r*(∞) was calculated from the average charge of the last 10 EPSCs in the train

Δ*t* is the stimulus interval in the train The RRP was estimated as RRP=*r*(1)/*fe*.

All currents were recorded using an Axopatch 200B amplifier, filtered (1 kHz) and digitised (10 kHz). Data were monitored online and analysed offline using WinEDR and WinWCP software (available free online at http://spider.science.strath.ac.uk/sipbs/software_ses.htm). Stimulus artefacts in representative traces were digitally removed for clarity.

## Statistical analysis

All experiments were performed at least three times. All calculations and graphs were performed with ImageJ, Microsoft Excel and GraphPad Prism software. P-values were calculated using two-tailed Student’s unpaired or paired (*in vitro* budding) t-tests, two-way ANOVA with repeated measures (electrophysiology), one-way ANOVA followed by Holm-Sidak correction for multiple comparison or one-way ANOVA for preselected pairs without correction for multiple comparison (clathrin biophysical properties). Detailed statistical information including statistical tests used, number of independent experiments, P-values and definition of error bars is described in individual figure legends.

## Additional methods

Details on methods for protein purification and assays, mouse genotyping and behavioural tests, electron microscopy and ultrastructure analysis are in SI Appendix, Materials and Methods.

## Data availability

Protein expression vectors, non-commercial antibodies and mouse strains produced in this study are available upon request. All data generated and analysed for this study and associated protocols are included in the main text or SI Appendix.

## ACKNOWLEDGMENTS

This work was supported by grants to F. M. B. from the Wellcome Trust (107858/Z/15/Z), to P.C.S. from the Medical Research Council (MR/M024083/1) and to P.C.S. and E. P. from Alzheimer’s Research UK (ARUK-PG2018A-002). L. R. was supported by a Wellcome Trust 4-year interdisciplinary PhD studentship. J. J. B. was supported by MRC funding to the MRC Laboratory of Molecular Cell Biology at UCL, award code MC_U12266B. P. N. D. was supported by a UCL Excellence Fellowship. The authors would like to acknowledge UCL IQPath, UCL Institute of Neurology, Queen Square London, WC1N 3BG for processing tissue slices for H&E staining and Massimo Signore (ICH) and Anna Crowley (KLB Transgenic facility, UCL) for the generation of the *Cltb*^ko^ mouse strain.

## AUTHOR CONTRIBUTIONS

The study was conceived by L. R. and F. M. B. with expert input from P. N. D., F. M., E. P. and P. C. S. and the project was managed by F.M.B. Electrophysiology experiments were conducted and analysed by F. M. All *in vitro* reconstitution, protein level and EM experiments and analyses were carried out by L. R. with expert assistance from J. J. B., F. M., E. P., P. N. D. and K. B. Mouse behaviour data and survival characteristics were collected by Y. C. and M. D. C. The paper was written by L. R., F. M. B., F. M. and P. C. S. and then all authors read and commented on the paper.

