## Supplementary material for "Clathrin light chain diversity regulates membrane deformation in vitro and synaptic vesicle formation in vivo": SI Appendix

**This file includes:**

Supplementary Methods

Supplementary Figures S1 to S7

Supplementary References

### SUPPLEMENTARY METHODS

#### Protein expression and purification

CLCs in pET28a were expressed with a thrombin-cleavable, N-terminal His-tag and purified by standard nickel affinity chromatography, followed by size exclusion chromatography on a Superdex 200 10/300 GL column (GE Healthcare). His-tags were removed from expressed CLC proteins by thrombin cleavage. His-tagged epsin1 in pET32c and His-tagged ΔENTH-epsin$^{144-575}$ in pQE32 were expressed and purified as previously described by standard nickel affinity chromatography and size exclusion chromatography on a Superdex 200 10/300 GL column (GE Healthcare). Hub and TDD fragments were expressed and purified as described (1, 2).

CCVs were purified from porcine brain as described previously (3) and clathrin triskelia purified by tris-extraction and size exclusion chromatography on a Superose 6 Increase 10/300 GL column (GE Healthcare). To generate CHC-only triskelia, endogenous CLCs were removed using NaSCN and separated from CHC by size exclusion chromatography on a Superose 6 Increase 10/300 GL column (GE Healthcare) (4).

For reconstitution of clathrin and Hub fragments with defined CLC compositions, CHC or Hub in buffer C (50 mM Tris-HCl pH 8.0, 50 mM NaCl, 2mM EDTA, 1mM DTT) at 0.8-1.0 mg/ml was incubated with purified CLCs at 0.7-1.0 mg/ml in in the same buffer or in 20 mM Tris-HCl pH 8.0, 200 mM NaCl at a 1:6.6 ratio (w/w) for 1 h on ice.

#### Electron microscopy

Freshly glow-discharged, homemade, carbon-coated formvar films on copper grids (Agar Scientific) were used for all EM applications. All specimens were observed using a Tecnai G2 electron microscope (Field Electron and Ion Company, FEI) at an acceleration voltage of 120 kV. Images were obtained using a SIS Morada digital camera and TIA software (FEI).

**Clathrin trimer stability**

Clathrin reconstituted with different CLC isoforms at a concentration of 1 mg/ml were dialysed overnight in 10 mM Tris pH 8.0 at 4°C. 2 µg of reconstituted clathrins were diluted in 10 mM Tris pH 8.0 containing 0%, 0.01%, 0.02%, 0.05%, 0.1%, 0.2% or 0.5% N-Lauroylsarcosine (SARC) to a final volume of 20 µl. After 5-10 min incubation on ice, SDS-free 4× loading dye (250 mM Tris pH 6.8, 40% glycerine, 0.02% bromophenol blue) was added to the samples and samples loaded onto an SDS-free 6% acrylamide gel to resolve mixtures of clathrin trimers and monomers by blue-native polyacrylamide gel electrophoresis (BN-PAGE). Two different running buffers, anode buffer (24.8 mM Tris-base, 192 mM glycine) and cathode buffer (24.8 mM Tris-base, 192 mM Glycine, 0.02% Coomassie Blue G250) were used for gel electrophoresis. All buffers were pre-chilled to 4°C and the gel chamber placed on ice during electrophoresis. Electrophoresis was carried out at 40 V for 15 min, then for 40 min at 90 V. Subsequently, the cathode buffer was exchanged for another cathode buffer of lower Coomassie concentration (24.8 mM Tris-base, 192 mM glycine, 0.002% Coomassie Blue G250) and electrophoresis continued for 2 h at 4°C. Samples were then transferred to nitrocellulose membrane and analysed by immunoblotting using TD.1 antibody (mouse monoclonal anti-CHC, made in house (5)) and X16 (mouse monoclonal anti-CLCa, made in house (6)) or CLTB antibody (rabbit antiserum against CLCb, Proteintech). Enhanced chemiluminescence substrate (ECL) was used to detect immunoblot signals with relevant secondary antibodies to measure the relative amounts of clathrin trimers and monomers within one sample using ImageJ software (NIH). Data from 3-5 repeats per clathrin reconstituted with one CLC isoform or CLC mixture were pooled and plotted against the logarithmic transform of detergent concentration and fitted using Prism 6 software (GraphPad).

**Cage size and lattice curvature determination**

Clathrin samples (native and CHC reconstituted with CLCs) were dialysed into assembly-promoting buffer A (100 mM MES, 1 mM EGTA, 0.5 mM $MgCl_2$, 2 mM $CaCl_2$). Cages formed from TDD and Hub fragments reconstituted with CLCs were assembled as described (1). Assembled cages were adsorbed to EM grids by placing grids on 10 µl droplets of samples at 0.2 mg/ml on Parafilm for 90 s. Excessive liquid was removed using filter paper and grids washed twice by transferring them sequentially onto two 15 µl droplets of buffer A. Samples were then stained with 2% uranyl acetate in water for 1 min and air-dried before subjected to EM analysis. Diameters of clathrin cages were measured from electron micrographs using ImageJ software (NIH). Distribution of diameters was averaged from the size distribution of 200 cages from three independent sets of experiments. For quantification, cage populations were grouped into cages smaller or larger than 90 nm in diameter and the proportions of the area under the curve for each population and experiment were averaged and tested for statistically significant differences between clathrin samples using Prism 6 (GraphPad).

**Flat lattice assembly and quality assessment**

For all incubations, EM grids were placed on 5-15 µl droplets on Parafilm at room temperature. Grids were firstly incubated with 0.04 mg/ml tag-free epsin1 or $H_6$-ΔENTH-epsin$^{144-575}$ in buffer G (25 mM HEPES pH 7.2, 125 mM potassium acetate, 5 mM magnesium acetate) for 30 min. Unbound protein was removed by transferring grids to two droplets of buffer G before incubating grids with clathrin samples at 0.05 mg/ml in buffer G containing 0.1% BSA for 30 min. After transferring grids to two droplets of buffer G, lattices were fixed with 3% glutaraldehyde in buffer G for 15 min and stained with 2% or 5% uranyl acetate in water for 1 min. For this, 2D Fast Fourier Transform (FFT) was produced using ImageJ software (NIH). The height of the peak corresponding to the periodicity of the lattice (at ~0.036 $nm^{-1}$) served as a measure for lattice quality – the higher the peak, the higher the quality of the lattice.

**Gold labelling of flat clathrin lattices**

EM grids were placed on 10 µl droplets of 0.04 mg/ml tag-free epsin1 in buffer G (25 mM HEPES pH 7.2, 125 mM potassium acetate, 5 mM magnesium acetate) and incubated for 30 min. Unbound protein was removed by two subsequent washes with buffer G before grids were incubated with clathrin at 0.05 mg/ml in buffer G containing 0.1% BSA for 30 min. After two further washes with buffer G, lattices were incubated for 10 min with a 1:5 dilution of 5 nm Ni-NTA-Nanogold® (Nanoprobes) in buffer G containing 0.1% BSA and washed with buffer G containing 25 mM imidazole for 1 min. Grids were then quickly rinsed with buffer G and lattices fixed with 3% glutaraldehyde in buffer G for 15 min. After two rinses with buffer A, lattices were stained with 2 or 5% uranyl acetate in water for 1 min.

**Staining of clathrin-coated liposomes for EM**

15 µl of the 15°C *in vitro* budding reaction (see Materials and Methods) were pelleted at 6000 g for 10 min in an Eppendorf 5424 R benchtop centrifuge (Eppendorf) and resuspended in ice-cold 3% glutaraldehyde in buffer G and fixed for overnight at 4 °C in suspension. Liposomes were then adsorbed to EM grids for 90 s. Specimens on grids were then stained for EM in the following sequence (7): 2× buffer A (quick rinse), 2× 0.1 M CaCo (rinse, 5 min), 1% $OsO_4$ in 0.1 M CaCo w/v (10 min), 2× 0.1 M CaCo (2× 2.5 min), deionized water (rinse), 1% tannic acid in water w/v (10 min), 2× deionized water (2× 5 min), 1% uranyl acetate in water w/v (10 min), 2× deionized water (2× 1 min).

**CLCb tissue expression analysis**

Tissue was cut into 1-2 mm pieces and lysed in RIPA buffer (50 mM Tris-HCl, pH7.4, 150 mM NaCl, 1% NP40, 0.5% Sodium deoxylcholate, 0.1% SDS and 1 mM EDTA) containing freshly added cOmplete, EDTA-free Proteinase inhibitor Cocktail (Roche). Tissue was further homogenised using a homogenizer (Polytron, PT 10-35) 3 times for 5 s and then centrifuged for 10 min at 14,000g at 4°C. The cleared lysate was then used for CLC expression analysis by standard SDS-PAGE and immunoblot procedures (as described in Materials and Methods for brain tissue).

**Genotyping of CLCb KO mice**

Mouse ear clip samples were incubated in 200 µl lysis buffer (100 mM Tris-HCl pH8.5, 5 mM EDTA, 0.2% SDS and 200 mM NaCl) containing 0.4 mg/ml proteinase K overnight at 55°C. The reaction was inactivated by incubating at 85°C for 40 min. PCR was performed using Phusion PCR Master Mix (NEB) using PCR instrument (Bio-Rad C1000T Thermal Cyclers). Primers (Sigma) are shown as below. Programme conditions were: 98°C for 10 min, [98°C for 30 s, Tm (melting temperature) for 30 s, 72°C for 30 s] (32 cycles), 72°C for 5 mins, and hold at 12°C. $T_m$ for the *Cltb* KO reaction was 60°C and 55°C for *Cltb* WT.

Primer sequences:

*Cltb* WT Fw: GCA GGA CAC TAG TGA CAA CC

*Cltb* KO Fw: TCA TTC TCA GTA TTG TTT TGC C

*Cltb* WT/KO Rv: TTC TAA GAC TAC ATC CTC ACC

**Electron microscopy of synapse ultrastructure**

Experiments were performed with mice aged 4-12-months. Female mice were used for ultrastructural experiments. Brain hemispheres were collected from homozygous KO mice and WT littermates and cut in sagittal orientation. Slices were fixed in 2% paraformaldehyde and 1.5% glutaraldehyde and stained sequentially with 1.5% potassium ferricyanide and 1% $OsO_4$, 1% thiocarbohydrazide, 2% OsO4, 1% uranyl acetate and 0.66% lead nitrate in aspartic acid. Samples were then gradually dehydrated and embedded for electron microscopy. The ultrastructure of excitatory synapses in four electron micrographs of the CA1 hippocampus perinuclear region per animal were analysed for the ultrastructure of excitatory synapses using ImageJ (NIH) and Prism 6 (GraphPad) software (n = 3 per genotype). To determine SV density, SVs in 300 nm vicinity to the centre of the PSD in each synapse were counted.

**Mouse behavioural tests**

All experiments were performed in accordance with the Animals Scientific Procedures Act UK (1986). Heterozygous *Cltb*$^{ko/+}$ mice were crossed to obtain homozygous *Cltb*$^{ko/ko}$ mice and WT littermates. Heterozygous Clta$^{ko/+}$ mice were crossed to obtain homozygous *Clta*$^{ko/ko}$ mice and WT littermates. Experiments were performed with mice aged 2-8-months. Male mice only were used for all behavioural experiments.

Rotarod latency (Accelerating Rotarod; Ugo-Basile 7650 model) was used to assess motor coordination. Mice were placed on the rotarod, which was slowly accelerated from 3 to 30 rounds per minute over 5 min. Mice were given 3 trials per day with inter-trial intervals of at least 10 min, over 2 days. Time to fall from the rotarod (latency) was recorded for each trial. Mice that remained on the rotarod for the whole 5 min trial were assigned a 300 s latency.

Grip strength (grip strength meter; Columbus Instruments) was used to measure muscle strength in the forepaws. The grip strength meter was positioned horizontally, and mice were held by the tail and allowed to grasp the metal pull-bar. The animals were then pulled back and the force applied to the bar and the moment the grasp was released was recorded. Mice were given 5 trials with 1 min inter-trial intervals and were tested over 2 days. The mean of results from 10 trials was assigned for the grip strength of each animal.

Grid-walking was used to assess spontaneous motor deficits and limb movements involved in precise stepping, coordination, and accurate paw placement. Adult mice were required to navigate over a wire mesh grid. After each trial, 70% ethanol was used to clean the apparatus. Mice were given 3 trials with 1 min inter-trial intervals and were tested over 2 days. Behaviour on the grid was recorded on camera and was analysed later by an investigator who was blind to the genotype. A foot-slip was scored when the paw completely missed a rung and the limb fell between the rungs. Counts of foot-slips for right forelimb, left forelimb, right hindlimb and left hindlimb were obtained. Total footsteps were also recorded. The mean of results from 10 trials was used for each animal.

**SUPPLEMENTARY FIGURES**

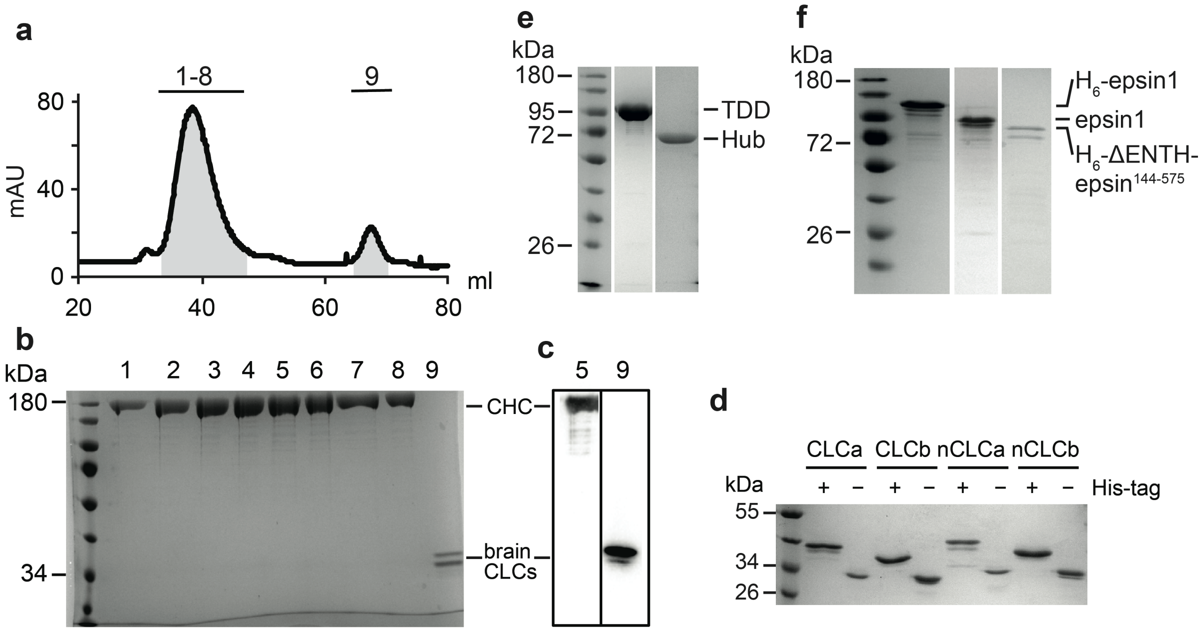


**Figure S1:** Protein purification from porcine brain tissue and recombinant expression. CCVs, isolated from porcine brain, were used as a clathrin source. **a** Size exclusion chromatography (absorbance at 280 nm in arbitrary units (mAU)) of native clathrin, purified from Tris-extracts of isolated CCVs, for separation of CHC triskelia and free endogenous CLCs after dissociation by thiocyanate treatment. Size exclusion fractions (millilitres (ml) of volume) as indicated (grey shading) were analysed by SDS-PAGE (bottom, lanes 1-8 and 9) and complete removal of CLCs confirmed by Coomassie staining (**b**) and immunoblot (**c**). TD.1 and CLTB antibodies were used for immunoblot analysis. **d** SDS-PAGE of recombinantly expressed CLCs with or without His-tags, which were removed by thrombin treatment as indicated (+/-). **e** SDS-PAGE (stained with Coomassie Blue) of recombinantly expressed Hub and TDD fragments, purified by standard nickel affinity purification and size exclusion chromatography. **f** SDS-PAGE (stained with Coomassie Blue) of recombinantly expressed full-length epsin1 constructs ($H_6$-epsin1 and tag-free epsin1 after thrombin treatment) and epsin1 fragment $H_6$-ΔENTH-epsin$^{144-575}$, purified by standard nickel affinity purification and size exclusion chromatography. The migration position of molecular mass markers is indicated in kilodaltons (kDa) at the left of immunoblots in b, e, f and g.

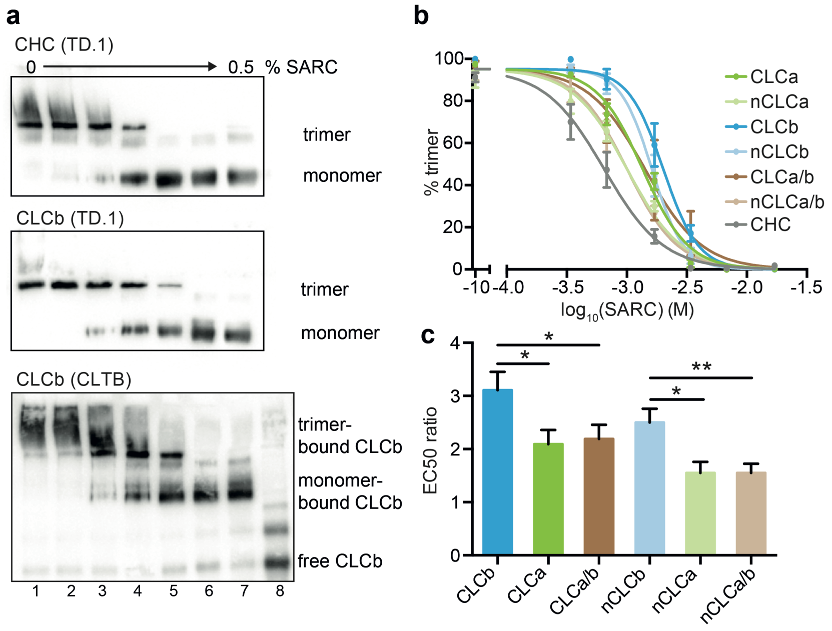


**Figure S2:** CLCs differentially stabilise the triskelion trimerisation domain. **a** Representative immunoblots for CLCb-reconstituted clathrin (CLCb) and CHC-only (CHC), which were incubated with increasing concentration (w/v) of N-lauryl sarcosine (SARC) detergent. Trimers and monomers were separated by native gel and proportions determined by immunoblotting using TD.1 antibody. The presence of CLCb bound to CHC (bottom panel, lane 1-7) was assessed by immunoblotting with CLTB antibody and compared to free CLCb (lane 8). **b** Dissociation profiles established as in **a** for clathrin reconstituted with indicated CLC isoforms, clathrin reconstituted with a 1:1 mix of CLCs or CHC alone ($3 < n < 7$) were fitted using an $EC_{50}$ shift model. **c** $EC_{50}$ ratio for dissociation from **b** (mean ± SEM, *$P < 0.05$, **$P < 0.01$, one-way ANOVA, $3 < n < 7$).

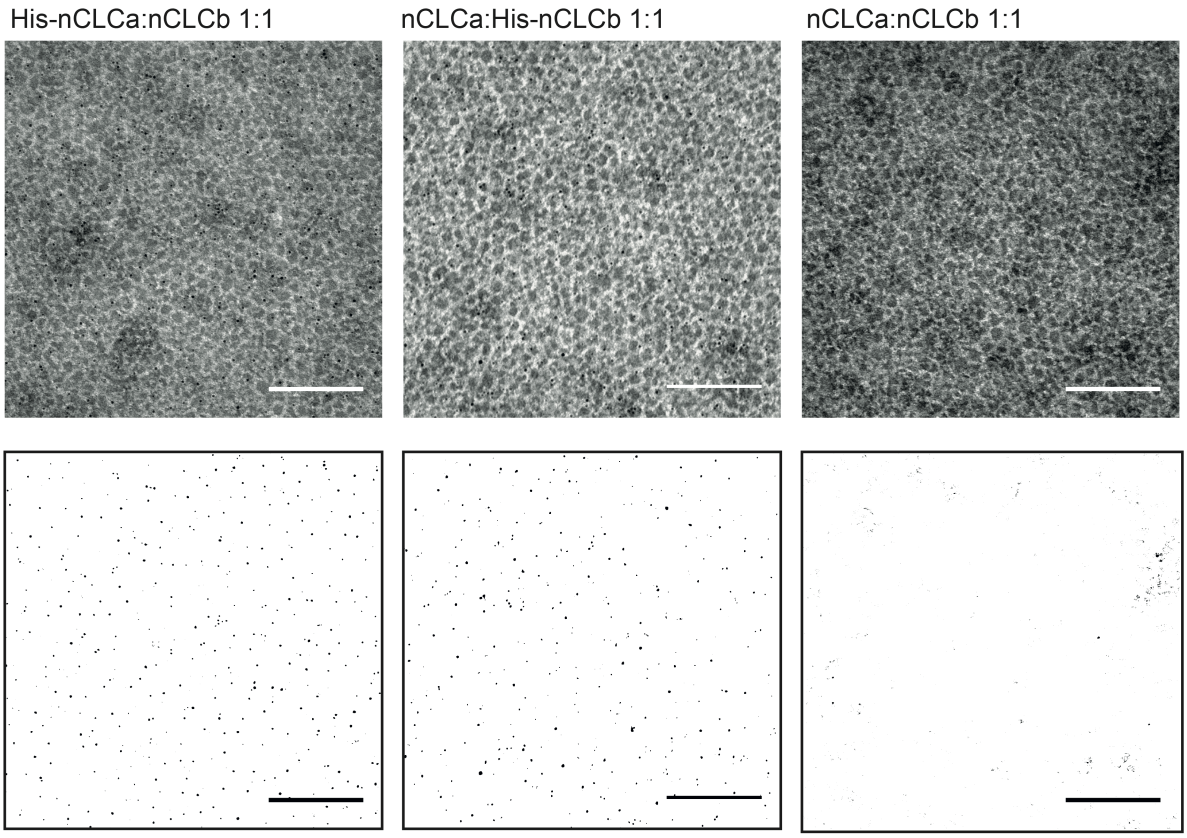


**Figure S3:** nCLCa clathrin and nCLCb clathrin co-assembly into flat lattices. EM images of lattices formed by co-assembly of clathrin reconstituted with the indicated CLCs (top) and thresholded images for visualisation of gold particles only (bottom). 1:1 mixtures of nCLCa clathrin and nCLCb clathrin, where either nCLCa, nCLCb or neither were His-tagged, were assembled on EM grids coated with tag-free, full-length epsin1 for lattice assembly. Incorporation of His-tagged triskelia was visualised by Ni-NTA-gold labelling. Scale bar: 200 nm.

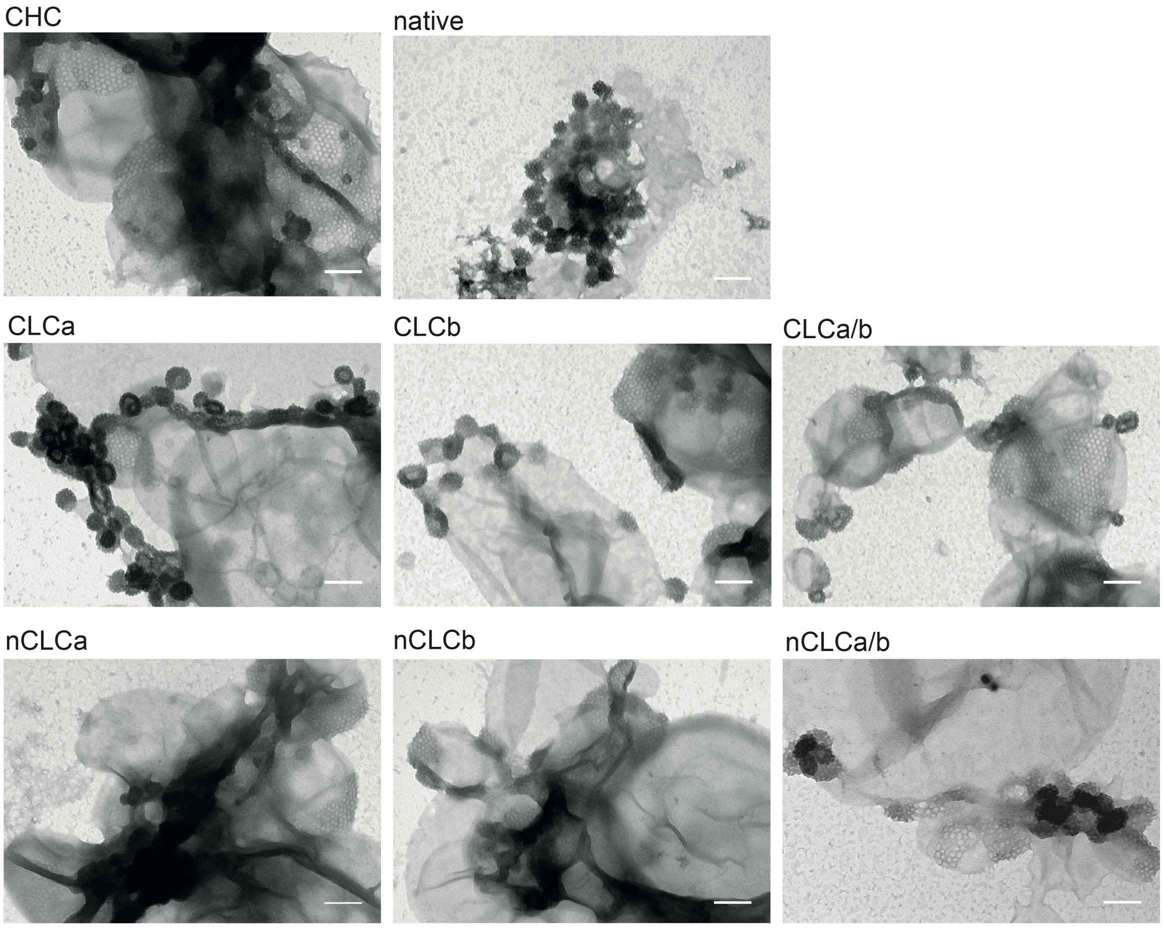


**Figure S4:** All reconstituted clathrins form intact lattices on liposome membrane *in vitro*. Liposomes made from brain polar lipid extracts containing 5% DGS-Ni-NTA lipids were bound with $H_6$-ΔENTH-epsin$^{144\text{-}575}$, chilled to 15°C and incubated with clathrin reconstituted with CLC isoforms as indicated (CLCa, nCLCa, CLCb, nCLCb), 1:1 mixtures of reconstituted clathrin (CLCa/b, nCLCa/b), CLC-free clathrin (CHC) or tissue-derived clathrin with native CLC composition (native) at 15°C for 30 min to allow lattice formation and vesicle budding. Samples were then fixed at 4°C overnight, adsorbed to EM grids and processed for visualisation of clathrin lattices by EM. Scale bar: 200 nm.

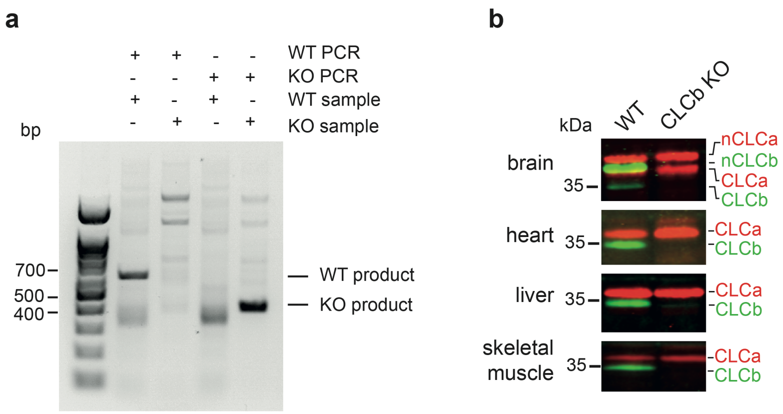


**Figure S5:** Confirmation of *Cltb* gene deletion and loss of CLCb protein in tissues from CLCb knockout (KO) mice. **a** Mouse genotyping by PCR. Mouse genotypes from tail tissue extracts of wild-type (WT) and CLCb KO samples were determined by two genotype-specific PCR reactions (WT PCR and KO PCR). PCR reactions produced products of around 600 base pairs (bp) for the WT and of 400 bp from KO genotype. Migration positions of base pair (bp) markers are shown at the left. **b** CLC expression in indicated tissues from homozygous CLCb KO and WT mice. Tissue extracts were prepared from brain, heart, liver and skeletal muscle tissue and (n)CLCa and (n)CLCb isoforms and splice variants were detected by immunoblotting with X16 (red) and CLTB (green) antibodies respectively. The migration position of molecular mass markers is indicated in kilodaltons (kDa) at the left of immunoblots.

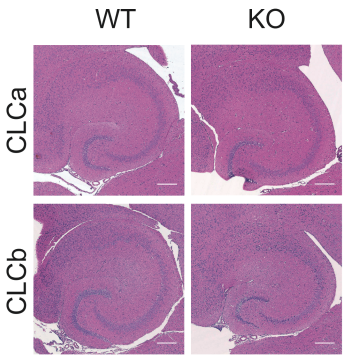


**Figure S6:** Hippocampus morphology in CLC KO animals and control mice. Representative H&E staining of sagittal sections from hippocampus of 9-10-month-old CLCa KO, CLCb KO and control animals (n = 4). Sectioning and H&E staining was performed by the UCL IQPath facility, Institute of Neurology.

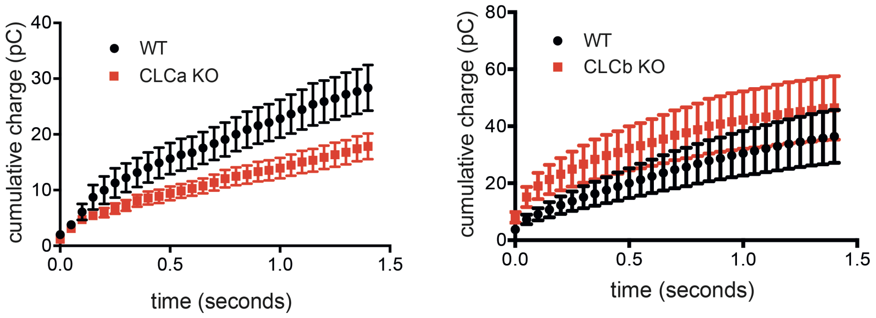


**Figure S7:** Cumulative charge plots for CLC KO and control mice. Plots show average cumulative charge (in picocoulomb (pC)) versus time (in seconds) during trains of stimulation at 20Hz for WT/CLCa KO mice (left) and WT/CLCb KO mice (right). These data are the basis for calculations of RRP size, fusion efficiency (fe) and vesicle recycling rate (α) in Fig. 4c and Fig. 4d, estimated with first order correction for vesicle recycling.
